# A novel resistance pathway for calcineurin inhibitors in the human pathogenic Mucorales *Mucor circinelloides*

**DOI:** 10.1101/834143

**Authors:** Sandeep Vellanki, R. Blake Billmyre, Alejandra Lorenzen, Micaela Campbell, Broderick Turner, Eun Young Huh, Joseph Heitman, Soo Chan Lee

## Abstract

Mucormycosis is an emerging lethal fungal infection in immunocompromised patients. *Mucor circinelloides* is a causal agent of mucormycosis and serves as a model system to understand genetics in Mucorales. Calcineurin is a conserved virulence factor in many pathogenic fungi and calcineurin inhibition or deletion of the calcineurin regulatory subunit (CnbR) in *Mucor* results in a shift from hyphal to yeast growth. We analyzed thirty-six calcineurin inhibitor resistant or bypass mutants that exhibited hyphal growth in the presence of calcineurin inhibitors or in the yeast-locked *cnbR*Δ mutant background without carrying any mutations in known calcineurin components. We found that a majority of the mutants had altered sequence in a gene, named here *bycA* (<u>by</u>pass of <u>c</u>alcineurin <u>A</u>). *bycA* encodes an amino acid permease. We verified that both *bycA*Δ, and the *bycA*Δ *cnbR*Δ double mutant are resistant to the calcineurin inhibitor FK506, thereby demonstrating a novel resistance mechanism against calcineurin inhibitors. We also found that the expression of *bycA* was significantly higher in the wild type strain treated with FK506 and in the *cnbR*Δ mutants, but significantly lower in the wild type without FK506. These findings suggest that *bycA* is a negative regulator of hyphal growth and/or a positive regulator of yeast growth in *Mucor* and calcineurin suppresses the *bycA* gene at the mRNA level to promote hyphal growth. BycA is involved in the *Mucor* hyphal-yeast transition as our data demonstrates a positive correlation between *bycA* expression, protein kinase A activity, and *Mucor* yeast-growth. Also calcineurin, independent of its role in morphogenesis, contributes to virulence traits including phagosome maturation blockade, host cell damages, and pro-angiogenic growth factor induction during interactions with hosts.

**Importance:** *Mucor* is intrinsically resistant to most known antifungals, which makes mucormycosis treatment challenging. Calcineurin is a serine/threonine phosphatase widely conserved across eukaryotes. When calcineurin function is inhibited in *Mucor*, growth shifts to a less-virulent yeast growth form which makes calcineurin an attractive target for development of new antifungal drugs. Previously we identified two distinct mechanisms through which *Mucor* can become resistant to calcineurin inhibitors involving Mendelian mutations in the gene for FKBP12, calcineurin A or B subunits and epimutations silencing the FKBP12 gene. Here, we identified a third novel mechanism where loss of function mutations in the amino acid permease encoding the *bycA* gene contribute to resistance against calcineurin inhibitors. When calcineurin activity is absent, BycA can activate PKA to promote yeast growth via a cAMP-independent pathway. Our data also shows that calcineurin activity, primarily contributes to host - pathogen interactions in the pathogenesis of *Mucor*.

## Introduction

Mucormycosis is a severe life-threatening infection caused by fungi belonging to the order Mucorales (1). People with weakened immune systems due to diabetes mellitus, neutropenia, or hematological and solid organ transplantation are at the highest risk of acquiring this infection (2, 3). It is the third most common invasive fungal infection in hematological and allogeneic stem transplantation patients following candidiasis and aspergillosis (1, 4). Over the past decades, there has been a global and ongoing rise in the incidence of mucormycosis, primarily due to the increasing number of diabetic patients and increased use of immunosuppressive drugs (5–10). Mucormycosis is also on the rise in immunocompetent individuals (11–14). Mucorales grow as molds in the environment and produce sporangiospores that can enter the host via inhalation resulting in pulmonary infection, through the skin due to trauma resulting in cutaneous infections, or through the nasal passages resulting in rhinocerebral infections (15–17). The spores can disseminate within the host, resulting in 95-100% mortality even with antifungal drug treatment (18). Mucorales are intrinsically resistant to most antifungals, which makes it very difficult to treat, and surgery is often required (19).

Calcineurin is a calcium-calmodulin-dependent phosphatase conserved widely across eukaryotes including pathogenic fungi (20, 21). Calcineurin is a heterodimer consisting of a catalytic and a regulatory subunit, and both subunits are required for calcineurin function. The role of calcineurin varies depending on the fungal species, for example in *Cryptococcus neoformans* and *Cryptococcus gattii* calcineurin is required for growth at high temperature (37°C) and at alkaline pH (22–24), while in *Candida* spp. calcineurin contributes to azole tolerance and is required for survival in serum, among other functions (20, 25). In *Aspergillus fumigatus*, calcineurin mutants exhibit delayed germination, hyphal growth with irregular branching, and abnormal septum (26). We have previously shown that in *Mucor* spp. calcineurin regulates dimorphism, in which the calcineurin inhibitor FK506 (tacrolimus) enforces *Mucor* to grow only as yeast (27).

Hyphal morphology is the predominant growth mode for *Mucor* spp., however by modulating respiratory conditions *Mucor* can be forced to grow as yeast as well (28). While aerobic conditions promote hyphal growth, low oxygen and high carbon dioxide conditions enforce yeast growth (29–32). Targeting components involved in mitochondrial or lipid metabolism can also promote yeast growth, even under aerobic conditions (33–36). In addition, previous studies have shown that the addition of cyclic AMP to *Mucor* in culture results in activation of cAMP-dependent kinase Protein Kinase A and promotes yeast growths (37–40). Wolff et al have also shown that during anaerobic yeast growth in *Mucor* an increased expression of PKA regulatory and catalytic subunits are exhibited than during aerobic hyphal growth (41). *Mucor* spp have four isoforms of PKA regulatory subunits, and each is differentially expressed depending on the growth conditions (38, 39). Calcineurin is involved in the genetic regulation of *Mucor* dimorphism as deletion of the gene encoding the regulatory subunit of calcineurin (CnbR) resulted in yeast-locked growth, even under aerobic conditions (27). *cnbR*Δ mutants are avirulent in a wax moth host model thereby making calcineurin an attractive target for antifungal treatment in mucormycosis (27). Also, mucormycosis incidence is low in patients receiving FK506 as an immunosuppressant (42). *cnbR*Δ mutants are also more susceptible to antifungal drugs such as Ambisome, micafungin, and posaconazole (43).

The cellular receptor for calcineurin is FKBP12, a member of the immunophilin protein family with cis-trans peptide prolyl isomerase activity (44). When FK506 is bound to FKBP12, it inhibits calcineurin phosphatase activity by binding to the interface between the calcineurin catalytic A and regulatory B subunits, thereby preventing substrates access to the active site (45, 46). FKBP12 also binds to rapamycin to inhibit the Tor pathway (47), and mutations in the FKBP12 gene confer resistance to both FK506 and rapamycin. Amino acid substitutions in the calcineurin regulatory B and catalytic A subunit surfaces that interact with the FKBP12-FK506 complex can also result in resistance to FK506 (48, 49). Another immunophilin, cyclophilin A (Cyp), serves as a cellular receptor for the drug cyclosporine A (CsA). When bound to Cyp, CsA inhibits calcineurin in a similar way to FKBP12-FK506 (50). Disruption of the gene encoding Cyp therefore confers resistance to CsA.

In our previous studies, calcineurin inhibitor-resistant *Mucor* strains, which exhibit hyphal growth instead of yeast growth, were found to have mutations in the FKBP12 gene or the calcineurin catalytic A or regulatory B subunit genes (27, 43, 51, 52). In addition, Calo et al., found that *Mucor* can also silence the FKBP12 gene to become transiently resistant to FK506 and rapamycin via an RNAi-dependent epimutation pathway (51, 52). In this study, we isolated mutants that do not follow the known calcineurin inhibitor resistance mechanisms. We identified a novel mechanism through which *Mucor* can become resistant to calcineurin inhibitors. We found that mutations or deletions in a novel gene, *bycA* (<u>by</u>pass of <u>c</u>alcineurin <u>A</u>), encoding an amino acid permease confers resistance to calcineurin inhibitors or loss of calcineurin regulatory B subunit. This gene has not been previously described in the calcineurin pathway, morphogenesis, or virulence in Mucorales. As a result, *bycA* mutation allowed us to separate the yeast – hyphal morphology switch from calcineurin function and demonstrate that calcineurin, independent of its function in regulating morphology, contributes to *Mucor* – host interactions.

## Results

### Isolation of calcineurin bypass mutants in the yeast-locked *cnbR***Δ** mutant background

The *cnbR*Δ mutant grows exclusively as a yeast (27). However, we isolated spontaneous mutants that exhibit hyphal growth in the *cnbR*Δ background. When the mutant (10^3^ cells per spot) was grown in solid YPD medium for a prolonged incubation at 30°C exceeding 5 days, we observed that hyphal sectors emerged from the yeast colonies (Fig S1A). After two rounds of single-streak dilutions, 17 independently isolated mutants (CnSp mutants) in the *cnbR*Δ background were isolated (Fig 1, Table 1, and Fig S1B). The hyphae of the mutants continued to grow, producing aerial hyphae decorated with sporangiophores containing sporangiospores (asexual spores, hereafter denoted spores). This finding shows that these mutants can complete the entire vegetative cycle without a functional calcineurin. We previously demonstrated that calcineurin function is required for hyphal growth (27). These spontaneous mutants however do not have a functional calcineurin but filament and produce spores. Thus, the mutants carry a genetic suppressor mutation(s) of the *cnbR*Δ mutation.

**Figure 1.**
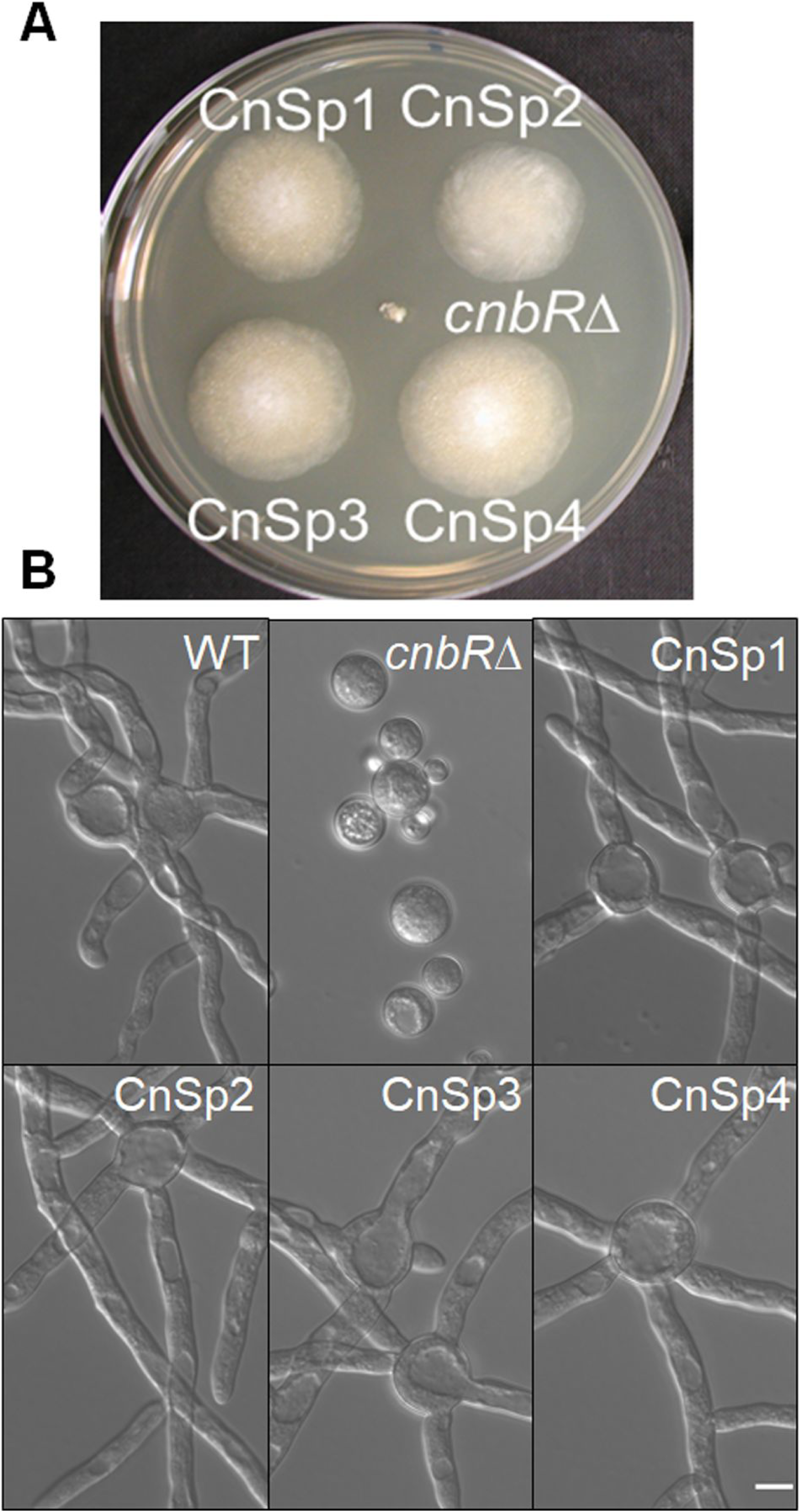
Calcineurin suppressor mutations (CnSp) in the *cnbR*Δ mutant background restore hyphal growth. A) Growth of calcineurin suppressor mutants (CnSp1 to CnSp4) and *cnbR*Δ mutant on a YPD agar plate at 30°C for four days post inoculation. While the *cnbR*Δ mutant shows smaller yeast colonies, the CnSp mutants form larger hyphal colonies. B) *Mucor* WT (R7B), *cnbR*Δ mutant, or CnSp mutants were grown overnight in liquid YPD medium with shaking at 30°C. Micrographs show that the CnSp mutants exhibit hyphal growth like WT (scale bar = 10 μm).

**Table 1.**
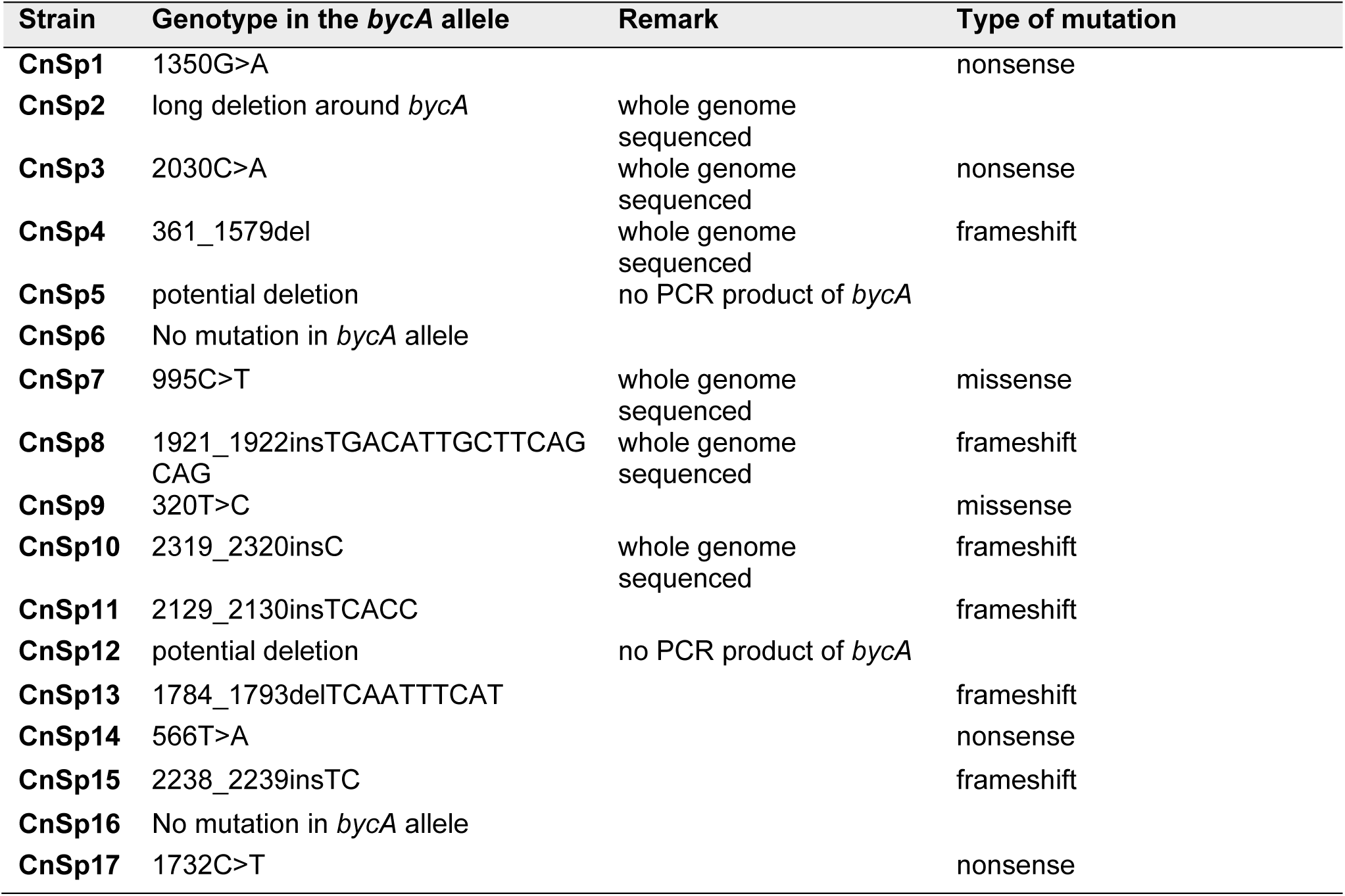
Characterization of the bypass mutants isolated in the *cnbR*Δ mutant background.

### Isolation of calcineurin bypass mutants in the *cnaB*Δ background

The calcineurin inhibitor CsA does not fully force *Mucor* to grow as yeast, unlike FK506. Instead, only when *Mucor* lacks the *cnaB* gene, one of the three calcineurin A catalytic subunit genes (*cnaA*, *cnaB*, and *cnaC*), does CsA fully inhibit hyphal growth and enforce a yeast-locked phenotype (43). We grew the *cnaB*Δ mutant on YPD medium containing 2 μg/ml of CsA, on which the mutant grew exclusively as yeast because calcineurin was inhibited by CsA. Similarly, after prolonged incubation on CsA medium, we observed that hyphal sectors emerged from yeast colonies as the strains became resistant to CsA (Fig S2). After two rounds of single-streak dilutions, we isolated 19 independently derived CsA-resistant (CSR) mutants (Table 2). The mutants exhibited hyphal growth in the presence of either CsA or FK506 (Fig 2).

**Figure 2.**
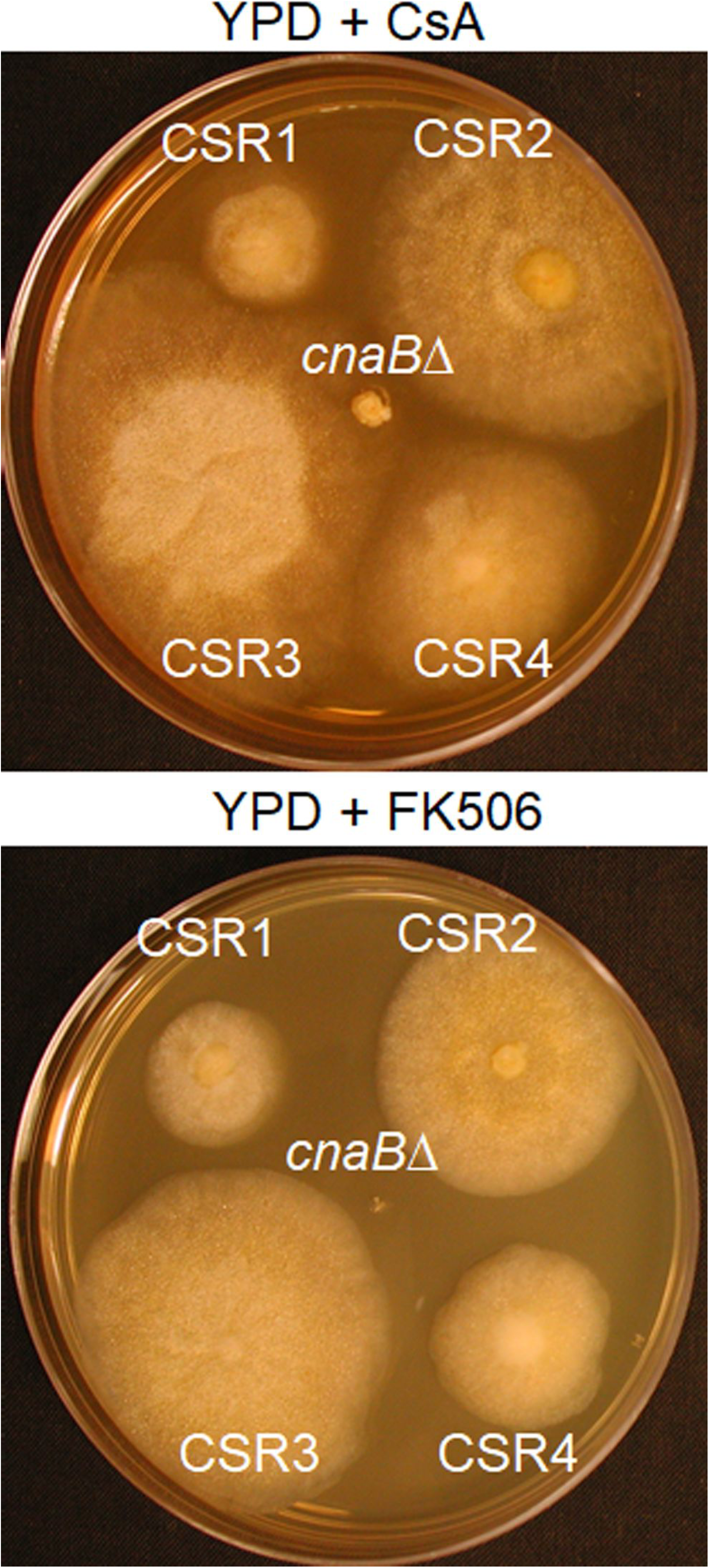
Calcineurin suppressor mutations (CSR) in the *cnaB*Δ mutant background confer resistance to calcineurin inhibitors. A) Growth of cyclosporine resistant mutants (CSR) exhibiting hyphal growth and *cnaB*Δ mutant exhibiting yeast growth when incubated on YPD agar with CsA (100 μg/ml) or FK506 (1 μg/ml) for four days at 30°C. CSR1, CSR2, CSR3, and CSR4 are shown. The other CSR mutants exhibited a similar CsA resistance phenotype (data not shown).

**Table 2.**
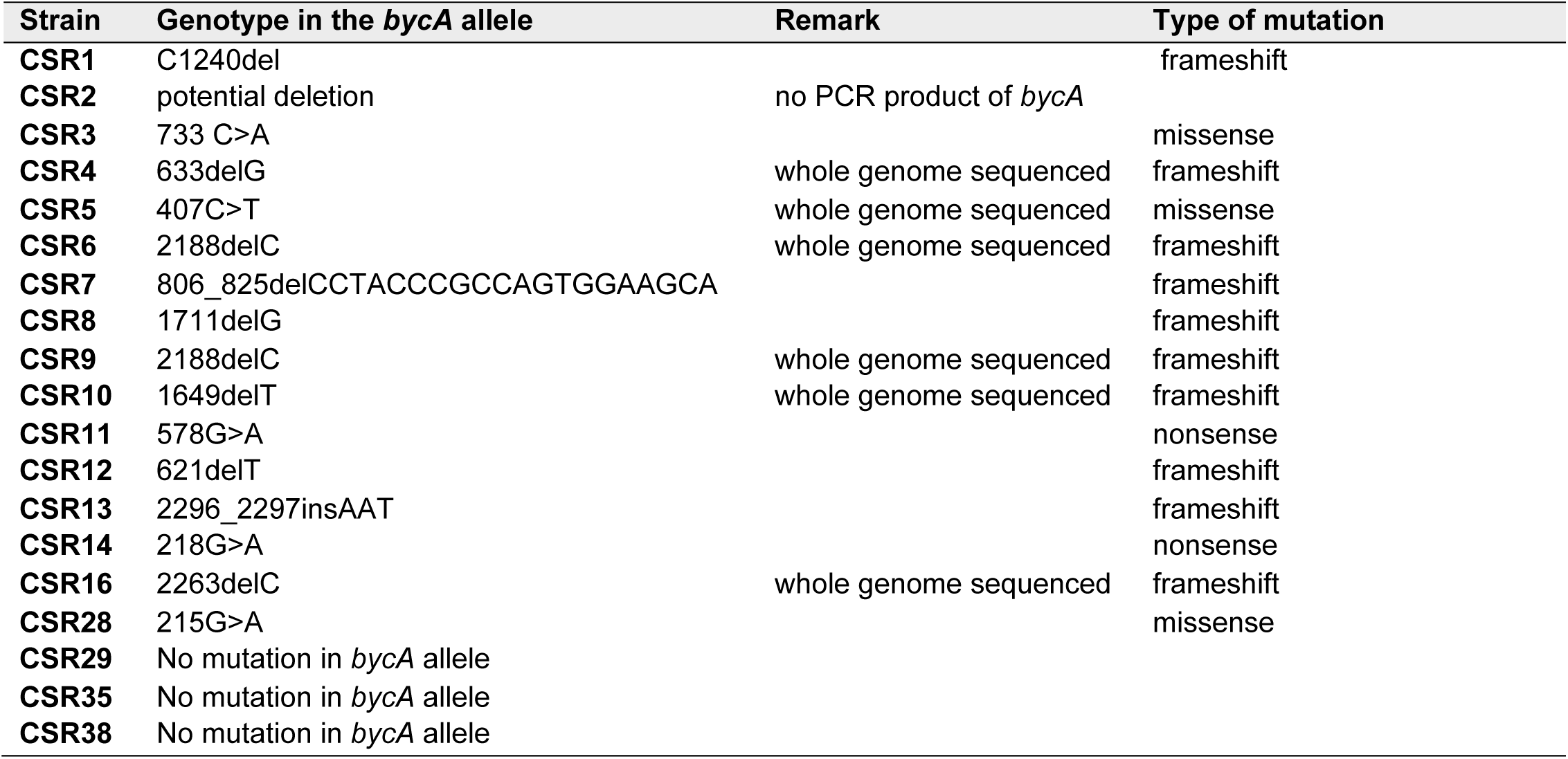
Characterization of the calcineurin bypass mutants isolated in the *cnaB*Δ mutant background.

Surprisingly, none of the mutants carried Mendelian mutations in genes for the calcineurin A or B subunit, FKBP12, or two subtypes of cyclophilin A (*cypA* and *cypB*) (data not shown). In addition, the mutants did not carry epimutations in the *cypA* and *cypB* genes, in which small RNA blots did not reveal any small RNAs derived from the *cypA* and *cypB* genes (51) (data not shown). Therefore, we hypothesized that the CSR mutants were also calcineurin bypass mutants that do not require calcineurin activity for hyphal growth.

### Spontaneous mutations in the *bycA* gene result in suppression phenotypes of the calcineurin mutation

To characterize the mutation(s) that results in bypass of the calcineurin requirement, we sequenced the whole genomes of 6 CnSp and 6 CSR mutant strains, along with the wild type strain MU402, which was used for transformation to obtain the *cnbR* and *cnaB* mutants, using the Illumina HiSeq platform. The whole genomes of each mutant strain were compared to that of the wild type MU402. Surprisingly, all twelve mutants carried DNA sequence modifications in a single common locus (Table 1 and Table 2). The modifications include an insertion of short sequences, single nucleotide polymorphisms, or deletions of a shorter or longer region in the locus. The gene in the locus was designated *bycA* (<u>by</u>pass of <u>c</u>alcineurin). We further sequenced the *bycA* gene in the remaining CSR and CnSp mutants and found mutations in the *bycA* gene in all but 5 mutants (Table 1 and Table 2). Thirty-one out of 36 suppressor mutants contained mutations in the *bycA* gene that could result in suppression of the calcineurin mutation.

### *bycA*Δ mutants are resistant to calcineurin inhibitors and *bycA*Δ *cnbR*Δ double mutants exhibit a hyphal morphology

We further verified that *bycA* is associated with the calcineurin pathway by generating *bycA* deletion mutants. The *bycA* gene in the wild type strain MU402 was replaced with the *pyrG-dpl237* marker (53) and gene replacement by recombination was confirmed by 5’ and 3’ junction PCR (Fig S3), ORF spanning PCR (Fig S3), and Southern blot (not shown). We also performed RT-PCR to confirm the *bycA* gene is not expressed in the mutants (see Fig 4A). Two independent mutants, MSL47.1 and MSL47.2 (*bycA*Δ::*pyrG-dpl237*), were obtained. The *bycA*Δ mutants exhibited hyphal growth and showed no growth defects (Fig 3A). On solid agar medium containing FK506, a *bycA*Δ mutant and CnSp4 produce a larger hyphal colony and aerial hyphae, whereas the wild type strain formed a smaller yeast colony (Fig 3B). These results indicate that *bycA*Δ mutants are resistant to FK506.

**Figure 3.**
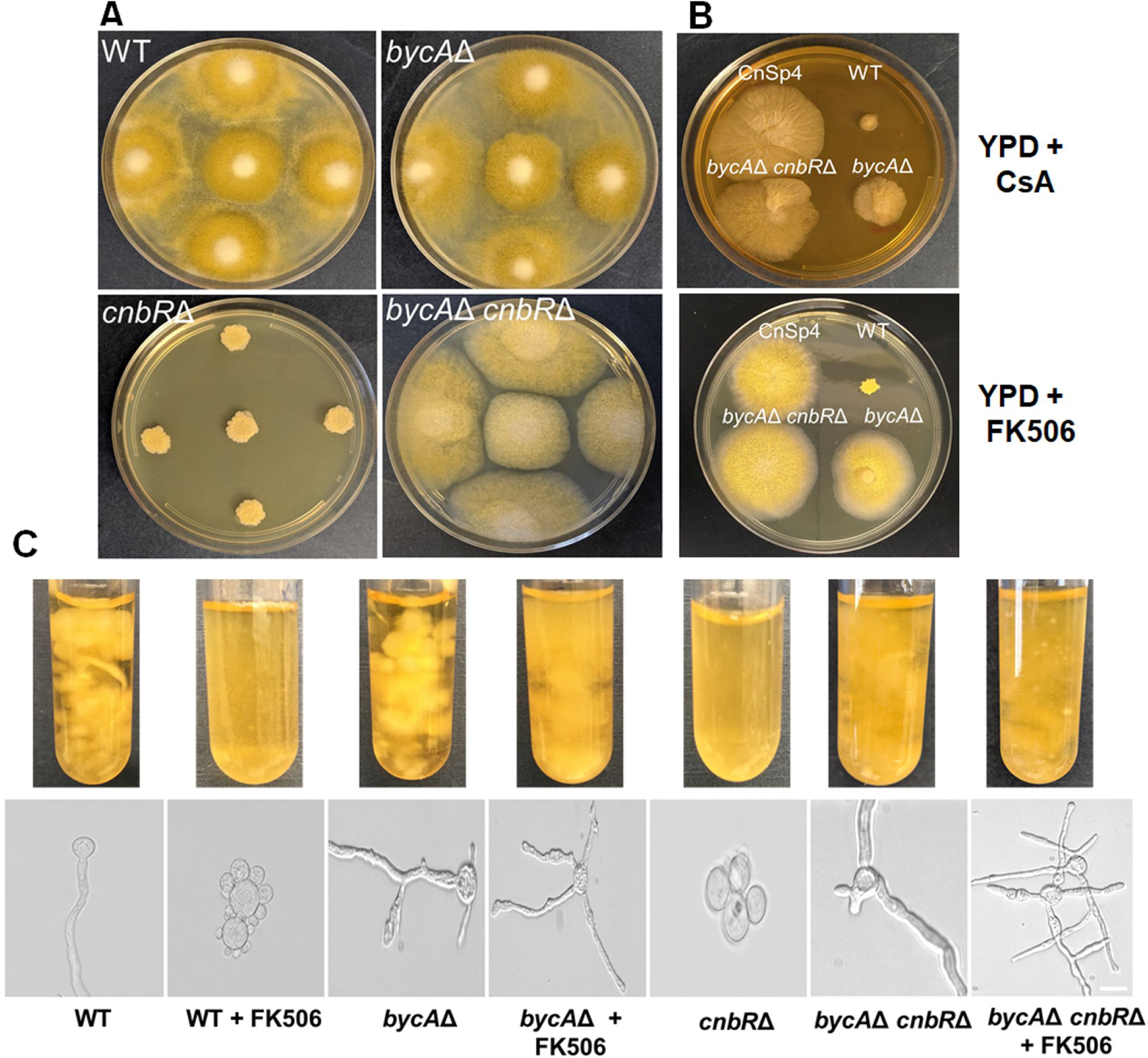
*bycA*Δ and *bycA*Δ *cnbR*Δ double mutant is resistant to calcineurin inhibitors. A) After three days of growth on solid YPD agar the *cnbR*Δ mutant grows as yeast; however, despite no calcineurin function, the *bycA*Δ *cnbR*Δ double mutant exhibits hyphal growth like WT and the *bycA*Δ mutant. B) In the presence of CsA (upper; 100 μg/ml) the *bycA*Δ *cnbR*Δ double mutant and CnSp4 formed larger hyphal colonies compared to WT, indicating resistance to CsA. No major difference in colony size was noted between *bycA*Δ mutant and WT. In the presence of FK506 (lower; 1 μg/ml) the WT forms a smaller yeast colony whereas the *bycA*Δ mutant, *bycA*Δ *cnbR*Δ double mutant and CnSp4 each form a larger hyphal colony. C) When *Mucor* was grown overnight in YPD medium containing FK506 (1 μg/ml) at 30°C with shaking, the *bycA*Δ mutant, *bycA*Δ *cnbR*Δ double mutant, and CnSp4 mutant exhibited resistance to FK506 as evident by larger biomass and hyphal morphology, whereas the WT is sensitive to FK506 as they not only form less biomass but also grow as a yeast. As expected, the *cnbR*Δ mutant remains in its yeast-locked form (scale bar = 20 μm).

**Figure 4.**
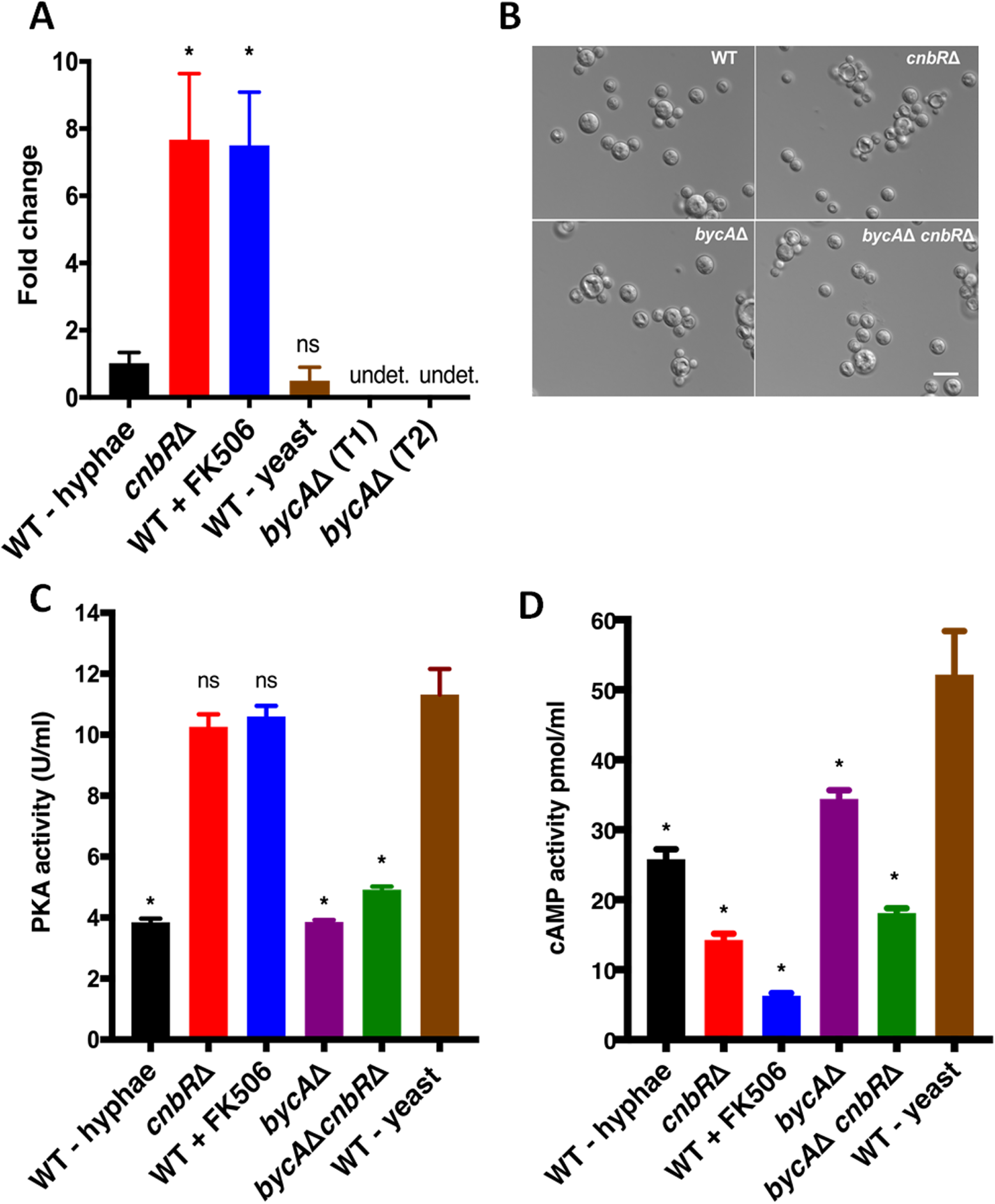
BycA is involved in the *Mucor* hyphal-yeast transition under aerobic conditions. A) Reverse transcriptase quantitative PCR showed that in the absence of calcineurin function (*cnbR*Δ and WT + FK506; yeast morphology) *bycA* expression was six-fold higher compared to WT (hyphal morphology) with calcineurin function suggesting that calcineurin regulates *bycA* expression at the mRNA level. When *Mucor* WT was grown anaerobically under high CO_2_ conditions (see methods) it grows as a yeast; however, there is no significant difference in *bycA* expression compared to WT-hyphae group. One-way ANOVA was significant (P = 0.0001). Dunnett’s post-hoc test was used to compare *cnbR*Δ, WT + FK506, WT-yeast with WT-hyphae group (*P<0.05). As expected, no expression of the *bycA* gene was detected in both of the *bycA*Δ mutants. B) When WT, *cnbR*Δ, *bycA*Δ, and *bycA*Δ *cnbR*Δ mutants were grown anaerobically overnight in high CO_2_ conditions, they all exhibited a yeast morphology thereby suggesting that neither calcineurin nor BycA has a role in anerobic morphological pathways (scale bar = 20 μm). C) Crude protein extracts (0.5 µg) were used to measure overall PKA activity, compared to WT-yeast (grown anaerobically) the WT-hyphae and *bycA*Δ have significantly lower PKA activity. In WT + FK506 and *cnbR*Δ mutant PKA activity remained higher, however *bycA*Δ *cnbR*Δ mutant has lower PKA activity despite no calcineurin function. One-way ANOVA was significant (P<0.0001). Dunnett’s multiple comparison test was used to compare each group with WT-yeast (*P<0.05). D) Crude extracts from 60 mg biomass were used to measure overall cAMP activity, compared to WT-yeast, the WT + FK506 and *cnbR*Δ had significantly lower cAMP activity thereby suggesting that under aerobic conditions, the PKA activity is elevated in *Mucor* yeast in a cAMP-independent manner. One-way ANOVA was significant (P <0.0001). Dunnett’s multiple comparison test was used to compare each group with WT-yeast (*P <0.05).

When grown on solid YPD media containing CsA (50 µg/ml), interestingly the *bycA*Δ mutants are not fully resistant to CsA, and instead exhibiting colony size comparable to wild type (R7B). However, CnSp4 exhibited larger colonies on media containing CsA (Fig 3B). It is possible that the CnaB catalytic A subunit is partially resistant to the inhibition by CsA, in which Cyp-CsA may not interfere with the function of CnaB-CnbR-calmodulin while still inhibiting CnaA-CnbR-Calmodulin and CnaC-CnbR-calmodulin. The *bycA*Δ mutants still harbor an intact *cnaB* gene and it is possible that partial inhibition of CnaB could result in a calcineurin independent phenotype. Therefore, *bycA*Δ mutants in the *cnaB*Δ background could exhibit full resistance against CsA, which is yet to be elucidated.

We further disrupted the *cnbR* gene in the *bycA*Δ background to generate *bycA*Δ *cnbR*Δ double mutants. The deletion and no expression of *cnbR* were confirmed by PCR and RT-PCR (Fig S4 and Fig S5), respectively, and Southern blot (data not shown). Two independent mutants MSL68.1 and MSL68.2 were generated (*bycA*Δ::*pyrG-dpl237 cnbR*Δ::*leuA*). As shown in Fig 3A, while *cnbR*Δ mutants grow exclusively as yeast, *bycA*Δ *cnbR*Δ double mutants exhibit filamentous growth like wild type and the *bycA*Δ mutant. *bycA*Δ *cnbR*Δ mutants are also completely resistant to either FK506 or CsA (Fig 3B), as they exhibit normal hyphal growth even in the presence of calcineurin inhibitors (Fig 3B and 3C). Taken together our results genetically validate our hypothesis that *bycA* mutations bypass the requirement of calcineurin for hyphal growth or suppress the lack of calcineurin.

### Calcineurin regulates BycA at the mRNA level

To determine a genetic link between calcineurin and BycA, the expression of *bycA* was examined under conditions in which calcineurin is either functional or non-functional. As shown in Fig 4A, compared to WT *bycA* gene expression was significantly higher in conditions in which calcineurin is not functional such as *cnbR*Δ mutant or the wild type in the presence of FK506. These results suggest that calcineurin negatively regulates the *bycA* gene at the mRNA level and BycA expression is positively correlated with *Mucor* yeast growth.

If grown in anaerobic (or microaerobic) conditions with high CO_2_, *Mucor* also grows as a yeast (54). Interestingly, under this condition, *bycA* expression remained low (Fig 4A). When *bycA*Δ or *bycA*Δ *cnbR*Δ mutants were grown anaerobically with high CO_2_ levels, they still grew as yeast (Fig 4B). These results suggest that BycA is involved in aerobic yeast growth but not in anaerobic yeast growth. Whether calcineurin inhibits the expression of the *bycA* gene transcriptionally or regulates the stability of the *bycA* mRNA remains to be elucidated.

### BycA serves as a link between calcineurin and protein kinase A

Previous studies by our groups and others have shown that addition of bicarbonate ions to *Mucor* culture, or by growth in high carbon dioxide conditions, is sufficient to induce yeast growth (27, 31), as the bicarbonate ions can activate adenylyl cyclase which results in the generation of cAMP and in turn activates cAMP-dependent kinase – protein kinase A (PKA) (55, 56) (Fig 8). Studies have also shown that there is an inverse relationship between PKA and hyphal morphology as the PKA regulatory subunit gene that inhibits the activity of PKA is more highly expressed during hyphal growth (41). Also, high levels of cAMP promote yeast growth, while low levels of cAMP are linked with hyphal morphology (28). Interestingly, even under aerobic conditions when *Mucor* was forced to grow as yeast by inhibiting calcineurin activity either genetically or by using FK506, the overall cellular PKA activity remained high (27). Therefore, PKA plays pivotal roles in yeast growth. Interestingly, in wild type when calcineurin is fully functional, the cellular PKA activity is low (27), suggesting an inverse correlation between PKA and calcineurin during morphogenesis. However, it was not clear how calcineurin and PKA are linked.

BycA is a putative amino acid permease and predicted to have ten transmembrane domains and one pectinesterase domain (Fig S6). As shown in Table 3, when *cnbR*Δ and *bycA*Δ *cnbR*Δ mutants were grown on YNB medium with either methionine, threonine, or arginine as a sole nitrogen source, the *cnbR*Δ mutants exhibited growth similar to growth on YNB complete medium, while *bycA*Δ *cnbR*Δ mutants did not produce hyphal mass. This shows that BycA is a bona fide amino acid permease. In *Saccharomyces cerevisiae* and *Candida albicans*, the general amino acid permease Gap1 is known to activate PKA in a cAMP-independent manner (57, 58). We hypothesized that in the absence of calcineurin activity, BycA may activate PKA to promote yeast-growth. To this end, we measured overall cellular PKA activity in the presence or absence of the *bycA* gene. As shown in Fig 4C, PKA activity was significantly higher in WT-yeast, *cnbR*Δ and WT + FK506 (when *bycA* expression was significantly higher) compared to WT - hyphae (when *bycA* expression is low). On the other hand, when the *bycA* gene is deleted, the *bycA*Δ or *bycA*Δ *cnbR*Δ mutants exhibited significantly lower cellular PKA activity compared to the conditions when the *bycA* gene is expressed. No significant differences were observed between WT - hyphae, *bycA*Δ, or *bycA*Δ *cnbR*Δ isolates.

**Table 3.** Summary for *cnbR*Δ and *bycA*Δ *cnbR*Δ double mutant growth in the presence of Methionine, Threonine, and Arginine as sole nitrogen source.

| Ammonium source | *cnbR*Δ mutant growth<sup>a</sup> | *bycA*Δ *cnbR*Δ mutant growth<sup>a</sup> |
| --- | --- | --- |
| None | - | - |
| Methionine | + | - |
| Threonine | + | - |
| Arginine | + | - |
| Casamino acids | ++ | ++ |

<sup>a</sup> + or ++ represent the extent of growth.

Interestingly, when we measured cAMP levels, compared to WT – yeast, the cAMP levels were significantly lower in *cnbR*Δ and WT + FK506 yeast groups despite high PKA activity (Fig 4D). This suggests that in aerobic conditions when calcineurin is absent, BycA increases PKA activity via a cAMP-independent pathway, which has yet to be elucidated.

### Phagosome maturation blockade upon phagocytosis by macrophages is dependent on calcineurin, not morphology

Phagocytosis followed by phagosome maturation or acidification in macrophages are essential innate immune pathways to control pathogens. Previously we have shown that when the macrophage cell line J774.A1 and primary macrophages are challenged with wild type spores or *cnbR*Δ yeast, they rapidly phagocytosed both spores and yeast, however only the macrophages with *cnbR*Δ yeast underwent phagosome maturation (43). This indicates *Mucor* spores escape innate immunity by blocking phagosome maturation as spores survive better than *cnbR*Δ yeast during co-culture with macrophages. However, it was not clear if the blockade of phagosome maturation by *Mucor* is dependent on its morphology or functional calcineurin because yeast cells lack calcineurin function. To address this question, we co-cultured WT spores, *cnbR*Δ yeast, and *bycA*Δ *cnbR*Δ double mutant spores with macrophages to monitor phagosome maturation. Macrophages containing *Mucor* spores or yeast cells were stained with Lysotracker Green DND-26 (Thermofisher). Lysotracker only stains acidic organelles in cells such as lysosomes or mature phagosomes and therefore can be used to determine whether phagosomes containing *Mucor* cells are acidic, an indication of phagosome maturation. As shown in Fig 5, only about 20% of macrophages challenged with either wild type (n=480) or *bycA*Δ (n=290) spores underwent phagosomal maturation, which is significantly lower when compared to macrophages challenged with *cnbR*Δ (n=279; yeast-locked). Interestingly, ∼80% phagosomes containing *bycA*Δ *cnbR*Δ double mutants (n=413) spores underwent maturation. Our data suggest a novel downstream function of calcineurin, independent of its function to govern morphology, is involved in the inhibition of phagosome maturation.

**Figure 5.**
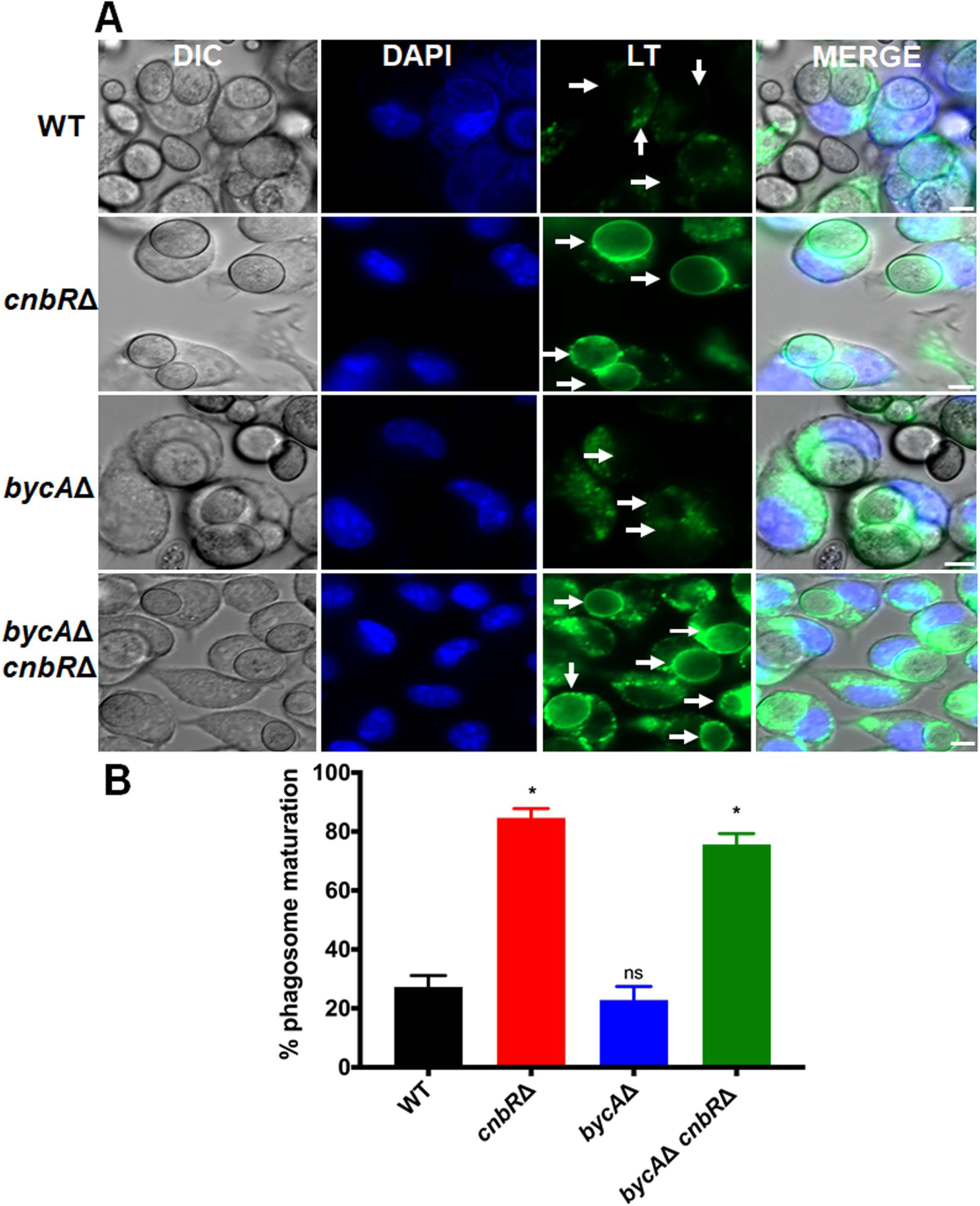
Phagosome maturation in macrophages containing *Mucor* is dependent on pathogen calcineurin function and not morphology. A) 5 x 10^5^ J774.A1 were challenged with *Mucor* spores or yeast at a MOI of 1 along with LysoTracker™ Green DND-26, and Hoechst 33342 stain (blue). White arrows indicate where phagosome maturation should be observed in the field (scale bar = 5 μm). B) Compared to WT, the macrophages containing *cnbR*Δ and *bycA*Δ *cnbR*Δ underwent significantly higher phagosome maturation. Data is shown as percentage maturation. The number (N) of macrophages containing *Mucor* counted for each group is: WT = 480; *bycA*Δ = 290; *cnbR*Δ = 279; *bycA*Δ *cnbR*Δ = 413. One-way ANOVA was significant (P <0.0001). Dunnett’s multiple comparison test was used to compare each group with WT (*P<0.05).

### *Mucor* without functional calcineurin causes reduced cell damage and induces less FGF-2 expression in endothelial cells

Mucormycosis is an angioinvasive disease, and hence Mucorales interaction and subsequent damage to the endothelium lining the blood vessels is an important step in disease pathology (59). We have previously shown that *Mucor* hyphae but not yeast (*cnbR*Δ) induces FGF-2 protein expression in lymphoblastoid cell lines and bone marrow macrophages (43, 60). We wanted to determine the extent to which calcineurin contributes to the endothelial damage and FGF-2 protein response by *Mucor.* To this end, we used Human Umbilical Vein Endothelial Cells (HUVECs) and challenged with wild type, *bycA*Δ, *cnbR*Δ or the *bycA*Δ *cnbR*Δ double mutant for a period of 24 hours. When compared to HUVECs challenged with wild type or *bycA*Δ spores, the *cnbR*Δ and *bycA*Δ *cnbR*Δ double mutant caused 70% less cytotoxicity in HUVECs as quantified by measuring LDH levels (Fig 6A) and induced significantly less FGF-2 protein production (Fig 6B). These data suggest a morphology independent function of calcineurin in regulating *Mucor* interaction with endothelial cells.

**Figure 6.**
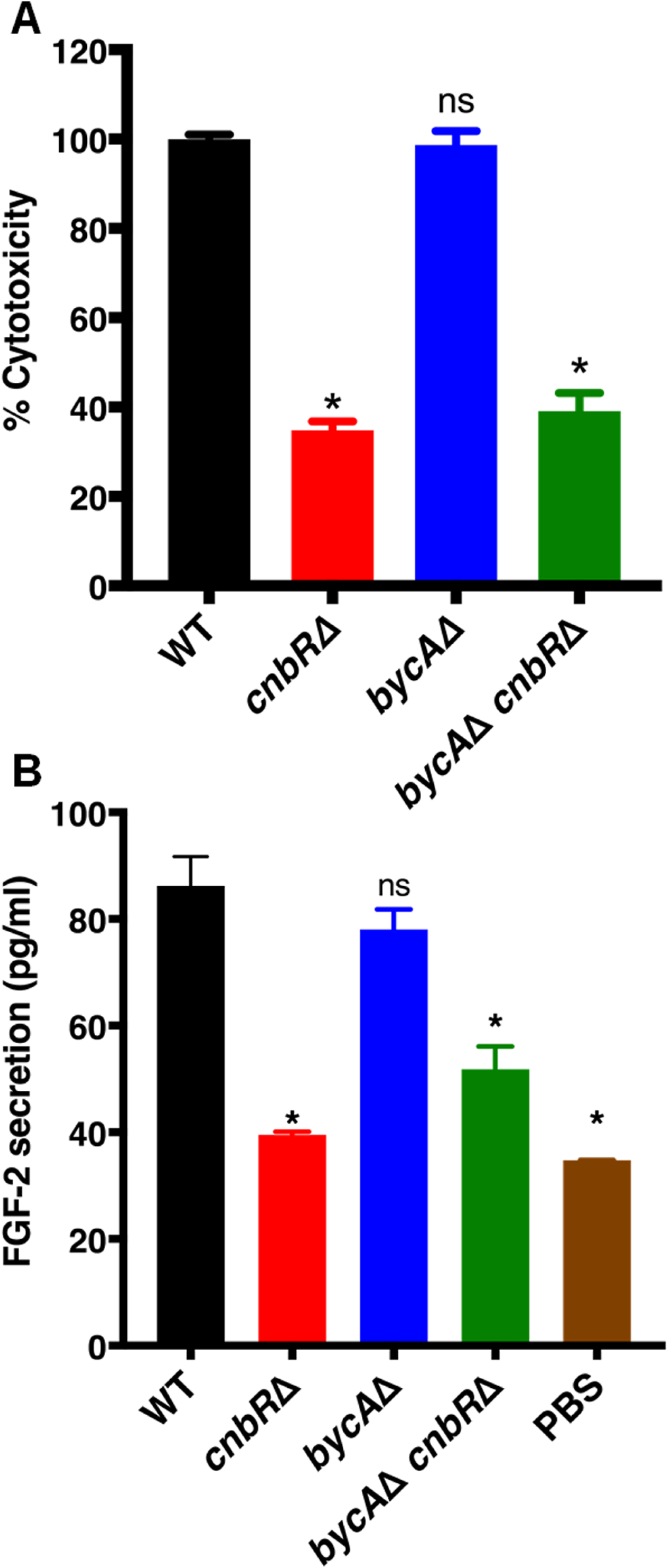
Calcineurin mutants cause less endothelial cell damage and FGF-2 protein secretion. 5 x 10^3^ HUVECs were challenged with *Mucor* spores or yeast at a MOI of 10. After 24 hours, the supernatant was collected. A) LDH levels were quantified which indicate cytotoxicity. Compared to WT, calcineurin mutants cause less damage. One-way ANOVA was significant (P <0.0001). Dunnett’s multiple comparison test was used to compare each group with WT (*P <0.05). B) FGF-2 levels were quantified using ELISA. The WT induce significantly higher FGF-2 protein secretion when compared to calcineurin mutants. One-way ANOVA was significant (P <0.0001). Dunnett’s multiple comparison test was used to compare each group with WT (*P <0.05).

### *bycA*Δ *cnbR*Δ double mutants are less virulent in a *Galleria mellonella* host model of mucormycosis

The yeast-locked *cnbR*Δ mutant is less virulent than wild type in a *G. mellonella* (wax moth) host (27). To test if calcineurin is involved in virulence, we injected wax moths (n=15/group) with 10,000 spores of either wild type, *bycA*Δ, *cnbR*Δ, the *bycA*Δ *cnbR*Δ double mutant, or PBS (mock) and survival was monitored for 7 days. As shown in Fig 7, all of the wax moth larvae challenged with wild type or *bycA*Δ spores succumbed to infection within a week, while *cnbR*Δ and *bycA*Δ *cnbR*Δ mutants exhibited a significantly higher survival rate. Interestingly, only about 65% of the wax moth infected with CnSp4 (spontaneous double mutant) or *bycA*Δ *cnbR*Δ survived, while 100% of the wax moth infected with *cnbR*Δ survived. We also injected wax moth larvae with 10,000 (1x), 20,000 (2x) or 30,000 spores (3x) of the *bycA*Δ *cnbR*Δ double mutant; as shown in Fig S7, it was only at 3x spore inoculum when 100% mortality was achieved.

**Figure 7.**
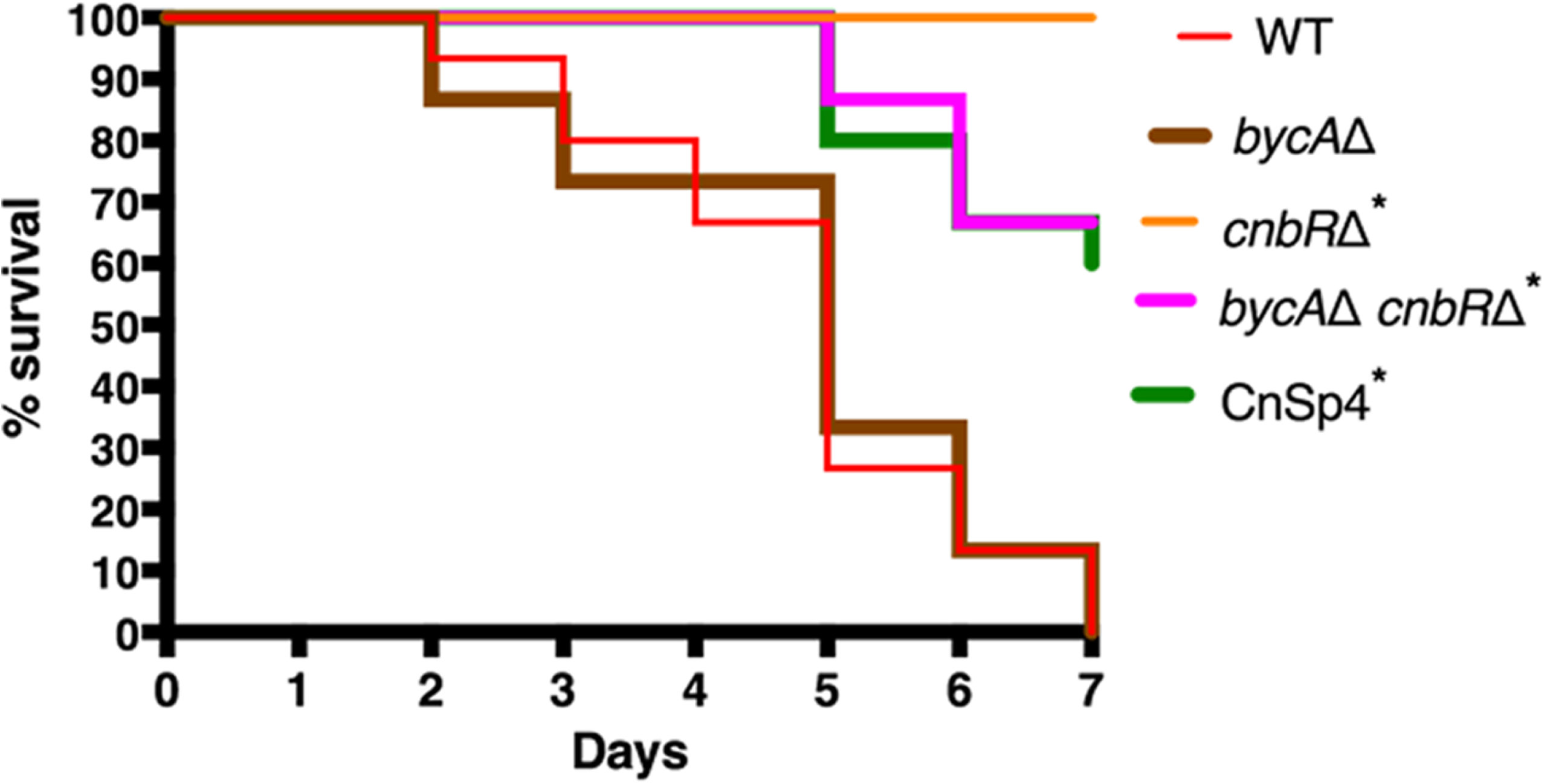
Calcineurin mutants are less virulent in a *Galleria mellanolla* (wax moth) model of mucormycosis. The wax moth larvae (n=15/group) were inoculated with 1 x 10^4^ *Mucor* spores or yeast in 2 μl PBS via injection into the last left proleg and were monitored for survival. All wax moth challenged with WT or *bycA*Δ succumbed to mortality within a week. The *cnbR*Δ was avirulent, whereas about 65% of wax moth inoculated with *bycA*Δ *cnbR*Δ or CnSp4 survived. A log-rank (Mantel-Cox) test was statistically significant (p <0.0001). A pair-wise comparison was also performed to compare the following groups: WT vs *bycA*Δ (p = 0.82); WT vs *cnbR*Δ or *bycA*Δ *cnbR*Δ or CnSp4 (*p <0.0001).

To determine virulence in an immunocompromised murine host, the wild type (R7B), *bycA*Δ, *cnbR*Δ, *bycA*Δ *cnbR*Δ double mutant, or PBS (mock) were inoculated via the intratracheal or intravenous routes. However, there was no significant difference in survival between the infected groups (Fig S8). It is interesting that calcineurin mutants are significantly less virulent in the wax moth larval host but are similarly virulent as WT in the murine model.

## Discussion

Calcineurin is conserved widely across pathogenic fungi and is involved in cell wall integrity, morphogenesis, and virulence (20, 21, 61). We have previously shown that when grown in the presence of a calcineurin inhibitor - FK506, *Mucor* exhibits yeast morphology even under aerobic conditions; however, when resistant to FK506 *Mucor* grows as hyphae (27). FKBP12 is the drug-receptor for FK506, and when bound, the FKBP12-FK506 complex inhibits the phosphatase activity of calcineurin by binding to the interface of the calcineurin regulatory and catalytic subunits (45). One of the common ways through which most pathogenic microbes become resistant to drugs is having mutations in the drug-receptor, in this case FKBP12 or catalytic A subunit or regulatory B subunits of calcineurin (27). Calo et al. have shown that *Mucor* can be transiently resistant to FK506 by triggering an RNA interference specific to the gene encoding FKBP12, resulting in silencing of the drug target gene (51, 52). In this study, we identified a novel resistance mechanism through which *Mucor* can also become resistant to calcineurin inhibitors. WGS and targeted sequencing of the genomes of spontaneous mutants (CnSp and CSR mutants) revealed DNA sequence alterations in the *bycA* locus in 31 out of 36 resistant isolates (Table 1 and Table 2).

### A novel link between calcineurin and BycA, an amino acid permease in *Mucor*

Based on our sequencing analysis we hypothesized that deletion of *bycA* should result in stable resistance to calcineurin inhibitors. Indeed, *bycA*Δ mutants are resistant to FK506. Unlike *cnbR*Δ mutants, which grow as yeast, the *bycA*Δ *cnbR*Δ mutant fully grow as hyphae like WT (Fig 3). This verifies that *Mucor* can become resistant to calcineurin inhibitors by having a second mutation in the *bycA* gene. *Mucor* has a secondary conserved mechanism that promotes hyphal growth even in the absence of a functional calcineurin. Interestingly, however, we did not observe Mendelian DNA mutations in the *bycA* locus in five of the mutants (out of 36) we performed sequencing on even though they exhibit the same phenotype as the remaining suppressors; one possibility is that *Mucor* may also produce small RNAi targeting *bycA* or there could be other mechanism(s) that confer *Mucor* resistance against calcineurin inhibitors. Future studies are required to elucidate these mechanisms further.

How is BycA linked to calcineurin and morphogenesis? It is not known if BycA is a direct post-translation modification target of calcineurin. However, our data suggest that calcineurin negatively regulates the expression of BycA at the mRNA level. It is also possible that calcineurin is involved in the stability of the *bycA* mRNA. Further investigation is ongoing to elucidate how calcineurin regulates expression of BycA. Interestingly, Chow et al revealed that, in *Cryptococcus neoformans*, a gene encoding a putative amino acid permease (CNAG_01118) is overexpressed when calcineurin is deleted and this is independent of Crz1, a well-known target of calcineurin (62). Whether deletion of the amino acid gene would result in suppression of calcineurin mutant phenotypes remains to be tested in *C. neoformans*.

### BycA as a missing link between calcineurin and PKA

Studies by us and others have shown that *Mucor* morphology is primarily dependent on: a) respiratory conditions and b) calcineurin and PKA activity (27–30, 37–39, 41). While aerobic conditions promote hyphal growth, *Mucor* grown in anaerobic conditions with low oxygen levels undergoes yeast growth. In anaerobic conditions with high CO_2_, *Mucor* grows as yeast, and has PKA activity increased via the bicarbonate-cAMP pathway (27, 28, 30). Previously we have shown that even under aerobic conditions when calcineurin function is inhibited by FK506 or absent in the *cnbR*Δ mutant, *Mucor* grows as a yeast and PKA activity remained high (27). The bicarbonate-cAMP pathway is likely not involved in aerobic conditions as the carbon dioxide levels are very low. It has been suggested that there is an antagonistic relationship between calcineurin and PKA in fungal systems in *Ustilago maydis* and *S. cerevisiae* (63, 64).

In this study, we found that calcineurin and PKA are inversely related through BycA, in which the overall PKA activity was lower in wild type hyphae with lower *bycA* expression, but higher in yeast with higher *bycA* expression (Fig 4A and 4C). PKA activity also remained lower in the *bycA*Δ *cnbR*Δ double mutant because even though calcineurin function is suppressed, the double mutant does not have BycA function to activate PKA and promote yeast growth. For the same reason, the *bycA*Δ *cnbR*Δ double mutants exhibit hyphal morphology. We also found that the amino acid permease BycA can activate PKA without a requirement for cAMP. Our finding that BycA can activate PKA is congruent with previous findings in *C. albicans* and *S. cerevisiae*, where the general amino acid permease Gap1 can activate PKA in a cAMP-independent manner (57, 58). It is possible that this is achieved through imbalance between the catalytic and regulatory subunits of PKA. In *S. cerevisiae*, it has been reported that kelch repeat homolog proteins 1 and 2 (Krh1 and Krh2) promote the association between PKA catalytic and regulatory subunits, and deletion of *krh1/2* leads to lower cAMP requirement for PKA activation (65). Since Krh proteins are evolutionarily conserved in eukaryotes, it is possible that such a mechanism may also exist in *Mucor.* Our future studies will further focus on determining how BycA activates PKA in *Mucor*.

Our previous studies showed that cellular PKA activity is elevated when calcineurin is inhibited in *C. neoformans* and *Rhizopus delemar* (27). How this link is achieved in these fungi remains be elucidated. In *C. neoformans*, there is evidence that higher expression of an amino acid permease is correlated with the phenotypes regulated by PKA; treatment with urea results in a 27-fold increase of an amino acid permease (CNAG_01118) and increased capsule production that is known to be regulated by PKA (66–68). Our current study investigates the links between calcineurin, amino acid permease, and PKA which could also exist in *C. neoformans* and possibly other fungi.

### Calcineurin governs virulence and plays important roles in host-pathogen interactions in *Mucor*

Studies from several pathogenic fungi have shown that morphology is linked with expression of virulence factors, colonization of the host, and evasion of host immune responses, etc (69, 70). The *cnbR*Δ mutants are significantly less virulent in a heterologous wax moth larva host model (27); however, it was not clear if the diminished virulence potential of *cnbR*Δ mutants was due to morphology, loss of the calcineurin, or both. In the current study using a hyphal growth strains lacking calcineurin - *bycA*Δ *cnbR*Δ double mutant, we identified a novel downstream function of calcineurin, independent of its function to govern morphology, that contributes to *Mucor* - host interactions and virulence. For example, macrophages challenged with mutants that lack calcineurin displayed significantly higher phagosome maturation compared to wild type, irrespective of their morphology (Fig 5). The ability of *Mucor* to cause damage to the endothelium was dependent upon calcineurin (Fig 6A).

Angiogenesis is the process by which new blood vessels arise from pre-formed blood vessels. Proteins such as Fibroblast Growth Factor-2 (FGF-2) and Vascular Endothelial Growth Factor (VEGF) promote angiogenesis (71). We have previously shown that the ability of *C. albicans* to induce FGF-2 is morphology-dependent, as non-filamentous strains of *C. albicans* fail to induce FGF-2 (72). We have also shown that in *Mucor* only WT, and not *cnbR*Δ mutants induce a host FGF-2 response (43). In this study we further found that both *bycA*Δ *cnbR*Δ double mutant also fails to induce FGF-2 (Fig 6B). Our data suggests that the ability of *Mucor* to induce host FGF-2 is dependent on a novel downstream function of calcineurin which is independent of its function to regulate morphology. Studies by us and others have shown that candidalysin from *C. albicans* and gliotoxins from *Aspergillus fumigatus* regulate the host FGF-2 response (72, 73). Hence, it is possible that toxins from *Mucor* (74, 75) also regulate host FGF-2 response. We are currently working on identifying the *Mucor* factor(s) that facilitates *Mucor* interactions with endothelial cells to induce FGF-2 response.

Larvae of *Galleria mellonella* have largely been used to study virulence and for testing antifungal drugs (27, 47, 74). The wax moth larvae possess an innate immune system that at the humoral and cellular levels is both structurally and functionally similar to mammals (76). We have previously shown that *cnbR*Δ mutants are avirulent in this host (27). In this study, we have shown that *bycA*Δ *cnbR*Δ double mutants are also less virulent (Fig 7). It is congruent that more wild-type spores survived compared to *cnbR*Δ yeast during interaction with bone marrow murine macrophages (Figure S9).

Interestingly, unlike in the wax moth host system, in the pulmonary and systemic murine model of mucormycosis, the *cnbR*Δ mutant also caused significant mortality like wild type. This observation may be due to the neutropenic host conditions caused by treatment with cyclophosphamide and cortisone. Calcineurin is required to escape innate immune cells by blocking phagosome maturation; however, lack or significant lower numbers of phagocytic cells resulted in higher mortality by the yeast-locked *cnbR*Δ mutant. In addition, treatment with cyclophosphamide could impact the phagocytic potential of the innate immune cells, as it was previously shown that treatment with cortisone acetate resulted in failure of alveolar macrophages to inhibit germination of *Rhizopus oryzae* (77). Alternatively, *Mucor* yeast is more immunogenic compared to *Mucor* spores and therefore the yeast might have caused mortality in a different manner than spores (43). This could be an analogy to the *Cryptococcus rim101* mutants (78). The *rim101* mutant lacks virulence traits and yet is more immunogenic compared to wild type; nevertheless, the mutants still cause mortality in a murine infection model via a substantial host immune response. Further study is required to determine if *Mucor cnbR*Δ yeast cause hyper immune responses in a murine lung infection model.

In summary, we identified a novel mechanism through which *Mucor* can become resistant to calcineurin inhibitors, where loss-of-function mutations in the *bycA* gene confer resistance against calcineurin inhibitors. This resistance mechanism is achieved by BycA regulating the activity of protein kinase A via a cAMP-independent pathway yet to be elucidated (Fig 8). As calcineurin is a major virulence factor in many pathogenic fungi, it is worth investigating if this relationship between calcineurin, BycA, and PKA is also conserved in other pathogenic fungal systems. Calcineurin also governs key *Mucor* – host interactions and is an attractive target for developing antifungals to treat mucormycosis.

**Figure 8.**
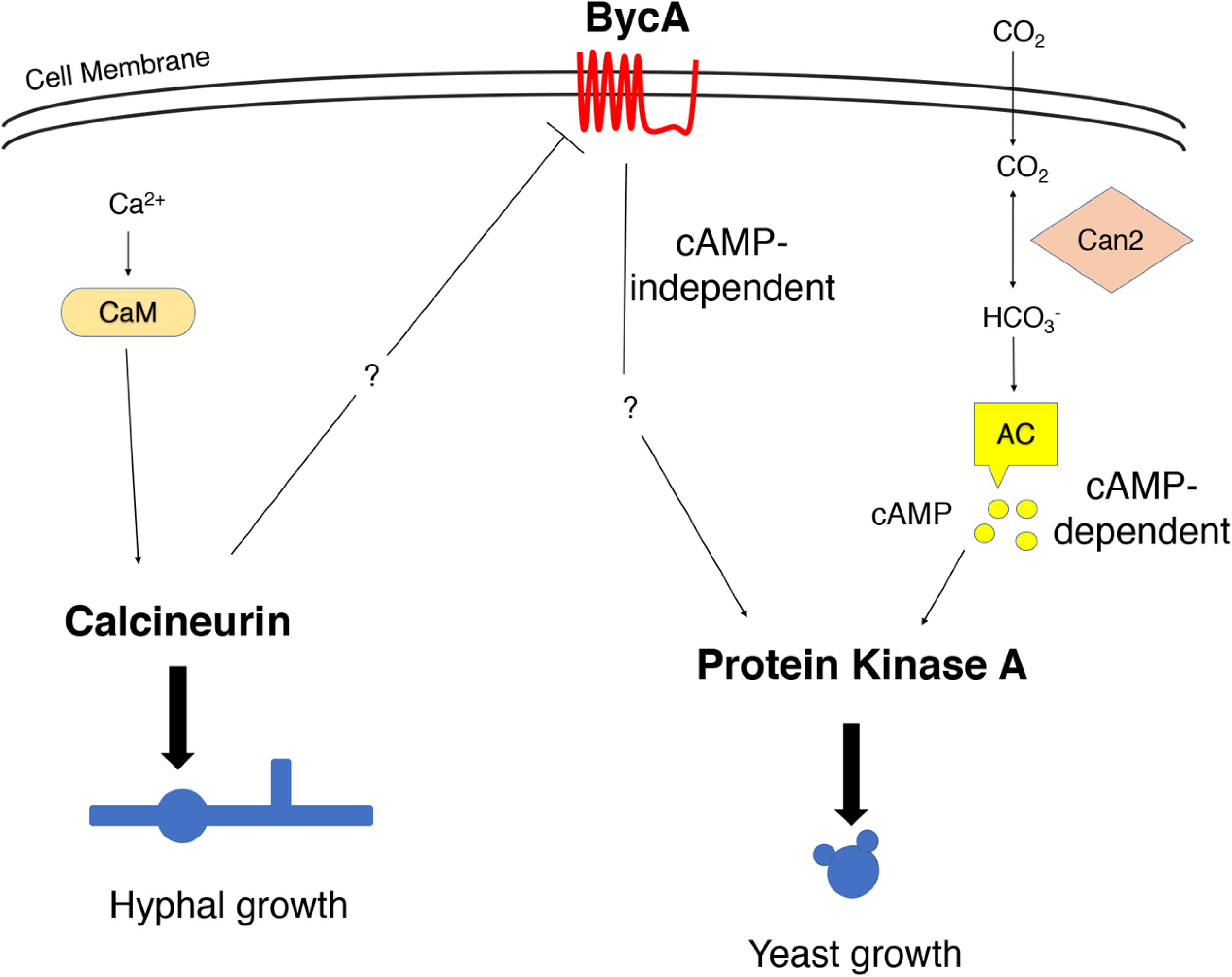
Calcineurin, BycA, and PKA in the morphogenesis of Mucor in aerobic conditions. Calcineurin is the master regulator of *Mucor* morphology. Active calcineurin positively regulates hyphal growth and negatively regulates yeast growth. This is achieved by suppressing the expression of the *bycA* gene thereby preventing an increase in PKA activity. Calcium-calmodulin activates calcineurin to promote hyphal growth under aerobic conditions. However, when calcineurin is not functional, *bycA* gene expression is significantly elevated. BycA then activates PKA in a cAMP-independent manner to promote yeast growth. In anaerobic conditions with high CO_2_ levels, PKA is activated through the CO_2_ – cAMP pathway. Neither BycA nor calcineurin has a defined role in the regulation of *Mucor* morphology under anaerobic conditions. Can2: carbonyl anhydrase, AC: adenylyl cyclase, and CaM: calmodulin.

## Materials and Methods

### Ethics Statement

All animal experiments were conducted at the University of Texas at San Antonio (UTSA) in accordance with the Institutional Animal Care and Use Committee (IACUC) guidelines and in full compliance with the United States Animal Welfare Act (Public Law 98-198) and National Institute of Health (NIH) guidelines. The animal protocol - MU104 used in this study was approved by the UTSA IACUC. The experiments were conducted in the Division of Laboratory Animal Resources (DLAR) facilities that are accredited by the Association for Assessment and Accreditation of Laboratory Animal Care (AAALAC).

### Fungal Strains and Growth Conditions

All fungal strains and plasmids used in this study are listed in Table S1A. For spore production, *Mucor* strains were inoculated and maintained on yeast peptone glucose (YPG, 3 g/L yeast extract, 10 g/L peptone, 20 g/L glucose, 2% agar, pH=4.5) or yeast peptone dextrose (YPD, 10 g/L yeast extract, 20 g/L peptone, 20 g/L glucose, 2% agar, pH=6.5) agar at 26°C or 30°C for four days. *cnbR*Δ mutants were maintained on YPD agar at 30°C. For in vitro, *Mucor –* host interaction studies or in vivo survival studies *cnbR*Δ mutants were grown overnight in liquid YPD at 30°C with shaking. For high CO_2_ conditions, the flasks were entirely filled with YPD broth, and the wild type (R7B) was inoculated on the bottom of the flasks. The flasks were sealed with parafilm and were left at room temperature overnight without disturbing. For testing with calcineurin inhibitors, *Mucor* strains were grown in liquid YPD or on YPD agar plates supplemented with FK506 (Astellas Pharma Inc.; 1 μg/ml) or CsA (LC laboratories; 2 or 100 μg/ml) at 30°C for 2 to 5 days.

### Generation of suppressor mutants CnSp and CSR

For generation of CnSp mutants, 10^3^ cells of *cnbR*Δ mutants were spotted on YPD agar and grown at 30°C for 5 or more days. Hyphal sectors emerging from yeast colonies were propagated to fresh YPD plates. For CSR mutant generation, *cnaB*Δ mutants were grown on YPD agar containing CsA (2 μg/ml) for 5 days or longer, and the hyphal colonies were propagated further.

### Whole Genome Sequencing and targeted sequencing of CnsP and CSR mutants

Genomic DNA of the 12 spontaneous mutant strains and MU402 strain was used for sequencing. A single end Truseq library was constructed with the genomic DNA and sequenced on the Ilumina HiSeq 2000 platform at the University of North Carolina at Chapel Hill School of Medicine. The reads were mapped to the genome of CBS277.49 (79) using the short-read component of BWA (80). SNP calling was performed using the Genome Analysis Toolkit (GATK ver. 2.4-9) pipeline and the Unified Genotyper with the haploid setting (81).

### Disruption of genes

The primers used in the study are listed in Table S1B. To disrupt *bycA* gene, nearly 1 kb upstream and downstream sequences flanking the *bycA* gene were PCR amplified (Phusion® High-Fidelity DNA Polymerase, NEB) using primers SL3 and SL4, and SL7 and SL8 respectively. The *pyrG*-*dpl237* marker (53) was amplified using SL5 and SL6 primers. All three fragments were gel excised, and 75 ng of each fragment was amplified using overlap PCR strategy with primers SL9 and SL10 such that the marker is placed between the flanking regions. The product was cloned into a TOPO vector (pSL26), which was then linearized using Sma1 enzyme and transformed into *Mucor* strain Mu402 (*pyrG^-^ leuA^-^*) via electroporation (82). The transformants were selected on minimal media with casamino acids pH=3.2 (MMC, 10 g casamino acids, 0.5 g yeast nitrogen base without amino acids and ammonium sulfate, 20 g glucose, 1 mg niacin, 1 mg thiamine, and 15 g agar in 1 L dH_2_O) containing 0.5 M sorbitol as described previously (82). Two independent transformants (MSL47.1 and MSL47.2; Table S1A) out of 12 were positive for disruption of *bycA*.

For disruption of the *cnbR* gene, 5’ and 3’ flanking regions were amplified using SL243 and, and SL281, and SL282 and SL244 respectively. A *leuA* marker was amplified using SCL737 and SCL738. Overlap PCR was performed using primers SCL286 and SCL287. The pSL58 plasmid containing the disruption cassette was linearized and used to transform MSL47.2. The transformants were selected on YNB medium (1.5 g ammonium sulfate, 1.5 g glutamic acid, 0.5 g yeast nitrogen base without amino acids and ammonium sulfate, 10 g glucose, 20 g agar in 1 L dH_2_O). Two out of 18 independent transformants (MSL68.1 and MSL68.2) were positive for disruption of *cnbR*.

### Reverse Transcriptase qPCR

Based on the ORF sequences, we designed primers for *bycA* (SL346 and SL347) and *cnbR* (SCL 578 and SCL 579) that span across two exons. *Mucor* strains were grown overnight in YPD liquid medium overnight, and the next day total RNA was isolated using TRIzol reagent according to the manufacturer’s instructions. 1 μg of RNA from each sample was used for cDNA synthesis (High Capacity cDNA Reverse Transcription Kit with RNase Inhibitor, Applied Biosystems), and qPCR was performed using SYBR Green QPCR Master mix (Thermo Scientific). Actin gene (primers SCL368 and SCL369) served as internal control. Three independent replications were performed for each experiment with RNA obtained from two independent preparations.

### Protein kinase A activity assay

*Mucor* strains were grown in appropriate conditions overnight and collection of crude protein extracts, and overall PKA activity measurement was performed using The DetectX® PKA (Protein Kinase A) Activity Kit according to the manufacturer’s instructions. Briefly, crude protein extracts were obtained by subjecting the samples to activated cell lysis buffer with protease inhibitor cocktail, PMSF, and sodium orthovanadate. Each sample was vigorously shaken in a bead beater (1 minute x 5 times with 1 minute cooling) followed by centrifugation to collect supernatants containing crude extracts. Total protein was quantified using the Bradford method (Biorad). 0.5 μg of each sample was used to determine overall PKA activity according to the manufacturer’s instructions.

### cAMP determination

*Mucor* strains were grown in appropriate conditions overnight and the next day cells corresponding to 60 mg (wet weight) were flash frozen and immersed in 0.1 M HCl. Crude extracts were obtained as described above. cAMP levels were measured according to the manufacturer’s instructions (Direct cAMP ELISA kit, Enzo).

### Phagosome maturation assay

Murine macrophage cell line J774.A1 was maintained in Dulbecco’s modified Eagle Medium (DMEM) with 10% FBS, and 200 U/ml mixture of penicillin and streptomycin antibiotics at 37°C, 5% CO_2_. Upon confluence, a cell scraper was used to detach cells, and 5 x 10^5^ cells were plated in each well of a 24 well glass plate. After 24 hours, *Mucor* spores or yeast were added at a multiplicity of infection of 1 along with 10 μM LysoTracker™ Green DND-26, and 1 μg/ml Hoechst 33342 stain. 30 minutes later a Zeiss Axio Observer microscope was used to image fields containing macrophages and *Mucor* cells. The experiment was performed in triplicate and was repeated on two different occasions.

### LDH assay and FGF-2 ELISA

Primary Human Umbilical Vein Endothelial cells (HUVECs) were purchased from Lonza and were maintained in Endothelial Basal Medium (EBM) containing hydrocortisone, ascorbic acid, Insulin Growth Factor, heparin, and FBS at 37°C, 5% CO_2_ according to the manufacturer’s instructions. Confluent cells were trypsinized and 5 x 10^3^ cells/well were seeded in a 96 well plate. After 24 hours, 5 x 10^4^ of appropriate fungal cells in PBS (MOI =10) or an equal volume of PBS were added to each well. 24 hours post infection, the supernatant was collected to quantify LDH levels (The CytoTox 96® Non-Radioactive Cytotoxicity Assay, Promega) or FGF-2 protein levels using ELISA (R&D systems). The LDH release was calculated as described previously (83).

### Virulence test

For production of *Mucor* spores, appropriate strains were grown on YPG agar at 26°C for 4 days under light. The *cnbR*Δ mutant was grown overnight in YPD broth at 30°C with aeration. Both the spores and yeast were washed two times with PBS, and different inoculums ranging from 1 x 10^4^ to 3 x 10^4^ cells in 2 μl PBS were injected into the wax moth host through the last left proleg. Differences between the survival curves were evaluated for significance using the Kaplan-Meier test. The experiment was performed on two different occasions with n = 15 animals for each group.

6 week old CD1 mice were immunocompromised with cyclophosphamide (250 mg/kg via the intraperitoneal route) and cortisone acetate (500 mg/kg via the subcutaneous route) every 5 days starting 2 days before inoculation (84). On the day of inoculation, the mice were anaesthetized using isoflurane and 1.5 x 10^6^ spores or yeast cells in PBS were introduced via the intratracheal route (pulmonary) were monitored routinely. Two independent experiments were performed with n=5 for each group. Data represented is from a single experiment.

### Statistics

Prism (Version 7, GraphPad Software Inc) was used to perform statistical analysis. A P value≤0.05 was considered significant.

## Acknowledgements

We are indebted to Jose Lopez-Ribot, Chiung-Yu Hung, Astrid Cardona, Stephen Saville, Ashraf Ibrahim, Floyd Wormley Jr, and Praveen Juvvadi for valuable discussions. We also thank Anna, Averette, Gloria Adedoyin, and Alexis Garcia for technical support for this study. This work was supported by NIH/NIAID R03 AI119617, a Korean Food Research Institution (KFRI) grant, and UTSA Research funds to S.C.L. S.C.L. holds a Voelcker Fund Young Investigator Pilot Award from the MAX AND MINNIE TOMERLIN VOELCKER FUND. J.H. was supported by NIH/NIAID R37 AI39115-21, R01 AI50113-15, and PO1 AI104533-05. J.H. is co-director and fellow of the Canadian Institute for Advanced Research Program Fungal Kingdom: Threats & Opportunities.

## Supporting information

**Figure S1. Emergence of hyphal sectors from *cnbR*Δ mutants, and isolation of calcineurin suppressor mutants.** A) *cnbR*Δ mutants were plated on YPD agar for 5 or more days at 30°C. B) The hyphae emerging from the yeast colonies were transferred to fresh YPD agar to propagate further. C) CnSP mutants produce hyphae decorated with sporangiophore containing sporangiospores like WT (scale bar = 20 µ).

**Figure S2. Emergence of hyphal sectors from *cnaB*Δ mutant grown on CsA containing medium.** *cnaB*Δ mutants were grown in the presence of CsA (2 μg/ml) for 5 or more days at 30°C. This resulted in emergence of hyphal sectors from yeast colonies.

**Figure S3. Confirmation of disruption of *bycA* gene by junction PCR and ORF spanning PCR.** A) Illustration of the *bycA*Δ::*pyrG*-*dpl237* and *bycA* alleles with ∼ 1 kb upstream and downstream flanking sequence. P1 (SL3) and P4 (SL8) recognize sequences outside the disruption cassette. P2 (SCL566) and P3 (SCL567) recognize *pyrG*-*dpl237*. P5 (SL182) and P6 (SL183) recognize *bycA*. B) 5’ end: P1 and P2 amplify a 2,203 bp region, and 3’ end: P3 and P4 amplify a 1,994 bp region in the *bycA*Δ::*pyrG*-*dpl237*, the primer pairs does not produce a fragment in the WT. P5 and P6 amplify 690 bp region within the *bycA* gene, so no amplification is noted in the mutants. Image not to scale.

**Figure S4. Confirmation of disruption of *cnbR* gene in the *bycA*Δ background by junction PCR and ORF spanning PCR.** A) Illustration of the *cnbR*Δ::*leuA* deletion in the *bycA*Δ::*pyrG*-*dpl237* strain and *cnbR* alleles with ∼ 1 kb upstream and downstream flanking sequence. P1 (SL243) and P4 (SL244) recognize sequences outside the disruption cassette. P2 (SL391) and P3 (SL392) are specific for *leuA*. P5 (SCL578) and P6 (SCL579) are specific for *cnbR*. B) 5’ end: P1 and P2 amplify a 2,388 bp region, and 3’ end: P3 and P4 amplify a 1,620 bp region in the *cnbR*Δ::*leuA* region, the primer pair does not produce a fragment in the WT. P1 and P4 produce a 5,711 bp product in the mutant and 3,342 bp product in the WT. P5 and P6 amplify 170 bp region within the *cnbR* gene, so no amplification is noted in the mutants. Image not to scale.

**Figure S5. Confirmation of disruption of *cnbR* gene in the *bycA*Δ background by RT-qPCR.** Primers SCL578 and SCL579 were used to perform qPCR on the cDNA obtained from WT and mutants (see methods). Actin (SCL368 and SCL369) served as a control. No expression for *cnbR* was detected in the *bycA*Δ *cnbR*Δ double mutant.

**Figure S6. Predicted structure of BycA showing ten transmembrane domains, and one cytosolic domain.** BycA is predicted to have 10 transmembrane domains, and one cytosolic domain. The transmembrane domains were predicted with the TMHMM 2.0 software package (http://www.cbs.dtu.dk/services/TMHMM/). The predicted cytosolic region contains a pectinesterase domain.

**Figure S7. *bycA*Δ *cnbR*Δ double mutant is lethal in a wax moth larva host only at higher inoculum.** The wax moth larvae (n=15/group) were inoculated with 1 x 10^4^ (1X) to 3 x 10^4^ (3X) *bycA*Δ *cnbR*Δ double mutant or WT spores (1X) in 2 μl PBS via the last left proleg and were monitored for survival. All wax moth challenged with WT or *bycA*Δ *cnbR*Δ (3X) succumbed to mortality within a week. 65% of wax moths inoculated with *bycA*Δ *cnbR*Δ (1X) and 45% of the wax moths inoculated with *bycA*Δ *cnbR*Δ (2X) survived. A log-rank (Mantel-Cox) test was statistically significant (p <0.0001). A pair-wise comparison was also performed to compare the following groups: WT vs *bycA*Δ *cnbR*Δ (1X) (*p <0.0001); WT vs *bycA*Δ *cnbR*Δ (2X) (*p =0.0014); WT vs *bycA*Δ *cnbR*Δ (3X) (p =0.9460).

**Figure S8. *Mucor* virulence in the murine model of systemic and pulmonary mucormycosis.** A) Cyclophosphamide and cortisone acetate treated CF1 mice were challenged with 1.5 x 10^6^ spores or yeast cells in PBS via lateral tail vein (systemic mucormycosis model). The animals (n = 5/group) were monitored for survival. A log-rank (Mantel-Cox) test was not statistically significant (p = 0.0585) suggesting no major difference in survival between WT and calcineurin mutant groups. B) For pulmonary mucormycosis model (n = 5/group), the same inocula were introduced via intratracheal route and monitored for survival. A log-rank (Mantel-Cox) test was not statistically significant (p = 0.1188).

**Figure S9.** Survival of wild-spores and *cnbR*Δ yeast during co-cultures with bone marrow macrophages. The spores survive the macrophages significantly higher compared to the yeast-locked mutant. The experiment was performed as described previously (Jung et al. 2019).

**Table S1** A) List of strains used in the study and B) List of primers used in the study.

